# Nuclear and structural dynamics during the establishment of a specialized effector-secreting cell by *Magnaporthe oryzae* in living rice cells

**DOI:** 10.1101/077693

**Authors:** Emma N. Shipman, Kiersun Jones, Cory B. Jenkinson, Dong Won Kim, Jie Zhu, Chang Hyun Khang

## Abstract

**Background:** To cause an economically important blast disease on rice, the filamentous fungus *Magnaporthe oryzae* forms a specialized infection structure, called an appressorium, to penetrate host cells. Once inside host cells, the fungus produces a filamentous primary hypha that differentiates into multicellular bulbous invasive hyphae (IH), which are surrounded by a host-derived membrane. These hyphae secrete cytoplasmic effectors that enter host cells presumably via the biotrophic interfacial complex (BIC). The first IH cell, also known as the side BIC-associated cell, is a specialized effector-secreting cell essential for a successful infection. This study aims to determine cellular processes that lead to the development of this effector-secreting first IH cell inside susceptible rice cells.

**Results:** Using live-cell confocal imaging, we determined a series of cellular events by which the appressorium gives rise to the first IH cell in live rice cells. The filamentous primary hypha extended from the appressorium and underwent asymmetric swelling at its apex. The single nucleus in the appressorium divided, and then one nucleus migrated into the swollen apex. Septation occurred in the filamentous region of the primary hypha, establishing the first IH cell. The tip BIC that was initially associated with the primary hypha becomes the side BIC on the swollen apex prior to nuclear division in the appressorium. The average distance between the early side BIC and the nearest nucleus in the appressorium was estimated to be more than 32 µm. These results suggest an unknown mechanism by which effectors that are expressed in the appressorium are transported through the primary hypha for their secretion to the distantly located BIC. When *M. oryzae* was inoculated on heat-killed rice cells, penetration proceeded as normal, but there was no differentiation of a bulbous IH cell, suggesting its specialization for establishment of biotrophic infection.

**Conclusions:** Our studies reveal cellular dynamics associated with the development of the effector-secreting first IH cell. Our data raise new mechanistic questions concerning hyphal differentiation in response to host environmental cues and effector trafficking from the appressorium to the BIC.

## Background

The filamentous ascomycete fungus *Magnaporthe oryzae* causes economically important blast disease on rice and other crops [1]. Several key cellular events that occur during rice infection have been documented (Fig. 1A). On the rice leaf surface, the fungus forms a specialized infection structure, called an appressorium, which produces a narrow penetration peg to breach the plant surface [2]. Upon penetration, the peg expands to form a filamentous primary hypha that grows into the rice cell and subsequently differentiates into bulbous invasive hyphae (IH) with pseudohyphal appearance due to regular constrictions at presumed septa [3, 4]. Cell cycle regulation is important for the appressorium development and IH proliferation [5, 6, 7], and semi-closed mitosis has been proposed to occur in the appressorium and IH [8]. The primary hypha and IH invade living host cells (biotrophy) while contained within a host-derived extra-invasive hyphal membrane (EIHM) [4, 9]. The IH grow throughout the first invaded cell before invading adjacent cells [4] (Fig. 1A). The IH colonize each new cell biotrophically, but eventually kill host cells (necrotrophy); the invaded cells lose viability by the time the IH move into adjacent cells [4, 10].

**Figure 1.**
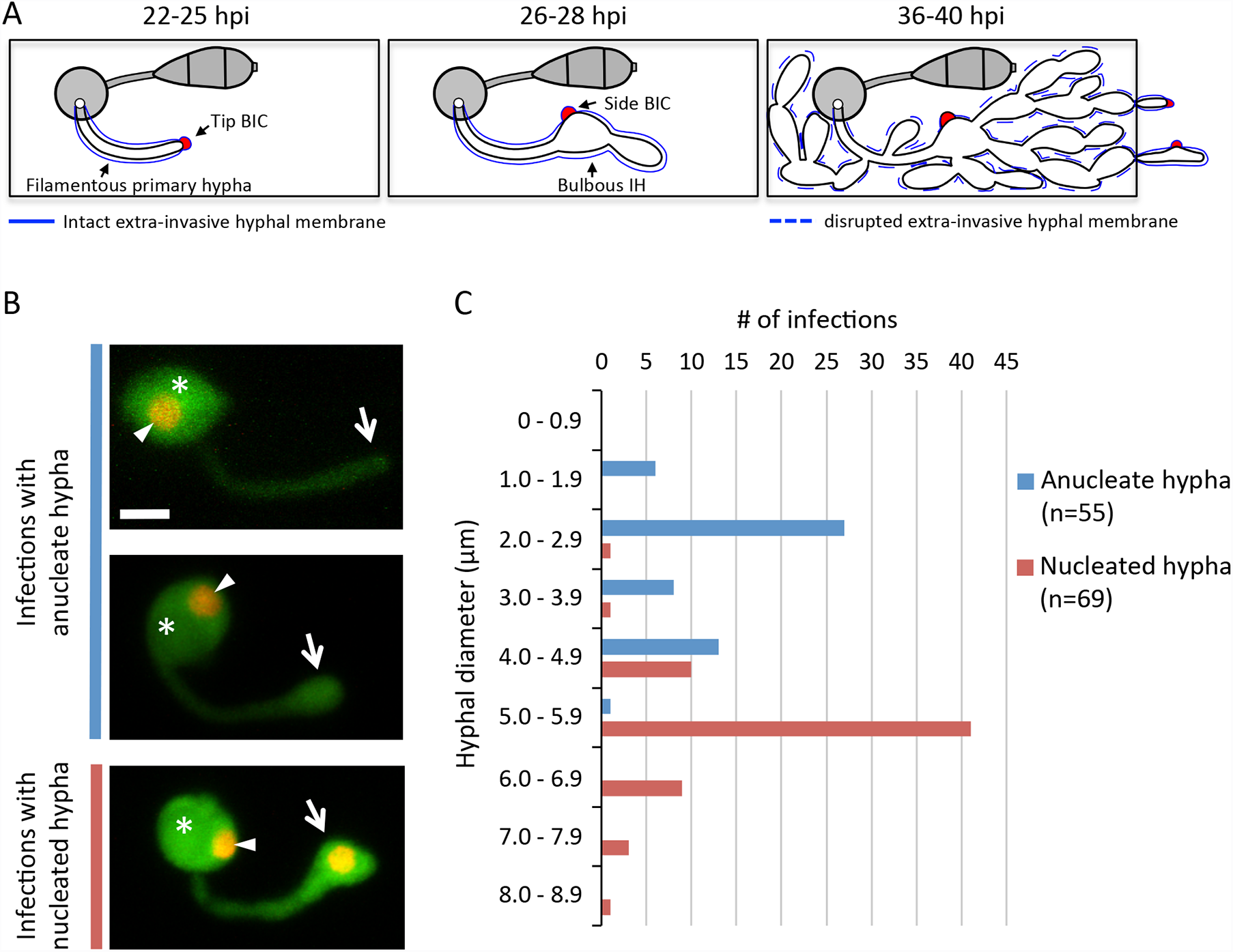
Primary hypha differentiates bulbous growth prior to mitosis. (A)Schematic diagram of IH development inside rice cells (rectangles), seen from above. Left (22-25 hours post inoculation; hpi): A filamentous primary hypha (white), derived from the appressorium (gray circle) on the rice cell surface, grows inside the rice cell. The hyphal apex is associated with the tip BIC (red). Middle (26-28 hpi): The primary hypha differentiates into bulbous IH, and the tip BIC becomes the side BIC (red). Right (36-40 hpi): IH have filled the first invaded rice cell and IH with secondary BICs are growing into a neighboring cell. 2 Hyphal development of *M. oryzae* CKF2138 in YT16 rice sheath cells at 26-27 hpi. Shown are maximum intensity projections of confocal optical sections (14 - 53 µm in total depth). Merged fluorescence shows fungal cytoplasmic EYFP (green) and nuclear tdTomato (red, but shown in yellow or orange due to the overlap of green and red). Top: The appressorium (asterisk) contained one nucleus (arrowhead) and produced an anucleate filamentous primary hypha. Middle: The tip of the primary hypha swelled but remained anucleate. Bottom: The bulbous end of the primary hypha had become nucleated. Arrows indicate the apical region of the primary hypha which correlates with the hyphal diameter measurements in (C). Bar = 5 µm. 3 Graph showing measurements of the diameter at the apical region of the primary hypha, corresponding to the location of arrows in (B), in CKF2138 infections at ~27 hpi. Groups of infection sites were plotted on the y-axis in 1 µm hyphal diameter increments and the x-axis shows number of infection sites (n=124). Anucleate hypha and nucleated hypha are differentiated by blue and red bars, respectively.

During the biotrophic invasion, the primary hypha and IH secrete cytoplasmic effector proteins (effectors; i.e., Pwl2, Bas1, Bas107, and AVRPiz-t) that enter the host cytoplasm and also apoplastic effectors (i.e., Bas4, Bas113, and Slp1) that are retained in the EIHM compartment [9, 11, 12, 13, 14, 15, 16]. These effectors are known or presumed to suppress host defense and facilitate infection [9, 13, 15]. Cytoplasmic effectors are preferentially localized in the biotrophic interfacial complex (BIC), which has been hypothesized as the site of effector translocation into the host cytoplasm [11, 17]. Two stages of BICs have been previously described [11] (Fig. 1A). The tip BIC is associated with the apex of the primary hypha. As the hypha develops, the BIC is positioned on the side of the first IH cell, and thus it is called the side BIC. The first IH cell, also known as the side BIC-associated cell, is a specialized effector-secreting cell essential for a successful infection. Giraldo et al (2013) have shown that this IH cell uses two distinct pathways for effector secretion. One is the conventional secretory pathway for secretion of apoplastic effectors into the EIHM compartment; the other is the exocyst components Exo70 and Sec5-involved pathway for secretion of cytoplasmic effectors into the BIC. In addition, this first IH cell is critical in its role as the mother cell of the subsequent biotrophic IH cells.

Although our knowledge of early biotrophic invasion has increased, there are questions that remain to be answered, such as how the first IH cell develops inside live host cells, what mechanisms control hyphal differentiation, and what cellular processes are involved in BIC development and effector trafficking. In this study, we provide insights into some of these questions by using live-cell imaging of *M. oryzae* invasion in susceptible rice cells. Our studies reveal a series of cellular events associated with the first IH cell development. These studies lead to new mechanistic questions concerning *in planta* hyphal differentiation and effector trafficking.

## Results and discussion

### Apical swelling of the primary hypha precedes mitosis

We generated *M. oryzae* transformants constitutively expressing EYFP (labeling the cytoplasm with green fluorescence) and histone H1-tdTomato fusion protein (H1-tdTomato; labeling nuclei with red fluorescence). These transformants exhibited wild-type morphology and pathogenicity. To characterize the development of the first IH cell upon host penetration, we used one of these transformants, CKF2138, and imaged 124 random infection sites at 22 - 28 hours post inoculation (hpi). Analysis of these images revealed three sequential growth stages (Fig. 1B): (1) the filamentous primary hypha with one nucleus in the appressorium, (2) the apically bulbous primary hypha with one nucleus in the appressorium, and (3) the apically bulbous primary hypha that contains a nucleus with another nucleus in the appressorium. We quantified these observations by measuring the diameter of the primary hypha close to the appressorium and the diameter at the apical region of the primary hypha before or after swelling (Fig. 1C). At all growth stages, the diameter of the primary hypha close to the appressorium was conserved with an average of 2.3 ± 0.4 µm (n=124). In infection sites with the anucleate primary hypha and a single nucleus present in the appressorium, the diameter of the primary hypha tip averaged 3.1 ± 1.0 µm (n=55). In infection sites displaying one nucleus in the appressorium and another one in the bulbous portion of the primary hypha, the diameter of the bulbous portion averaged 5.6 ± 0.9 µm (n=69), frequently ranging from 4.0 to 6.9 µm. Taken together, these data indicate that nuclear division and migration typically occur when the apical tip of the primary hypha has swollen over 4 µm in diameter.

### Mitosis in the appressorium and subsequent migration of a single nucleus into the first bulbous IH cell

To determine nuclear dynamics during development of the first IH cell, we conducted time-lapse microscopy of rice cells invaded by *M. oryzae* mitotic reporter strain CKF1962 expressing H1-tdTomato and GFP-NLS [8]. After plant penetration, nuclear division occurred in the appressorium, and subsequently one mitotic nucleus migrated through the primary hypha to arrive in the swollen portion of the primary hypha, while the other nucleus remained in the appressorium (Fig. 2A; n=14). The timing of nuclear division was consistent with our prediction that nuclear division occurs in infections that had a swollen primary hypha tip lacking a nucleus (Fig. 1C). Intriguingly, the nucleus that remained in the appressorium appeared to associate with the penetration pore as the other nucleus migrated through the primary hypha to the swollen hypha tip (Fig. 2A; n=19). Our observation of nuclear division within the appressorium contrasts a previous report, in which the appressorial nucleus migrates to the primary hypha and then undergoes mitosis [5]. We considered the possibility that the discrepancy might have risen because different strains of *M. oryzae* and rice were used, specifically *M. oryzae* O-137-derived strain (H1-tdTomato) infecting rice strain YT16 in our study and *M. oryzae* Guy11-derived strain (H1-tdTomato) infecting rice strain CO-39 in the other study [5]. However, we ruled out this possibility because we observed the consistent pattern of nucleus division within the appressorium followed by migration of a single nucleus into the primary hypha when *M. oryzae* Guy11-derived strain (H1-tdTomato) infects YT16 (Fig. 2B; n=4) or CO-39 (Fig. 2C; n=3). The reason for this discrepancy remains to be determined. It is worth noting, however, that our findings are consistent with *Colletotrichum gloeosporioides*, in which nuclear division occurs in the appressorium, and then one nucleus migrates into the penetration hypha, whereas the other nucleus remains in the appressorium [18].

**Figure 2.**
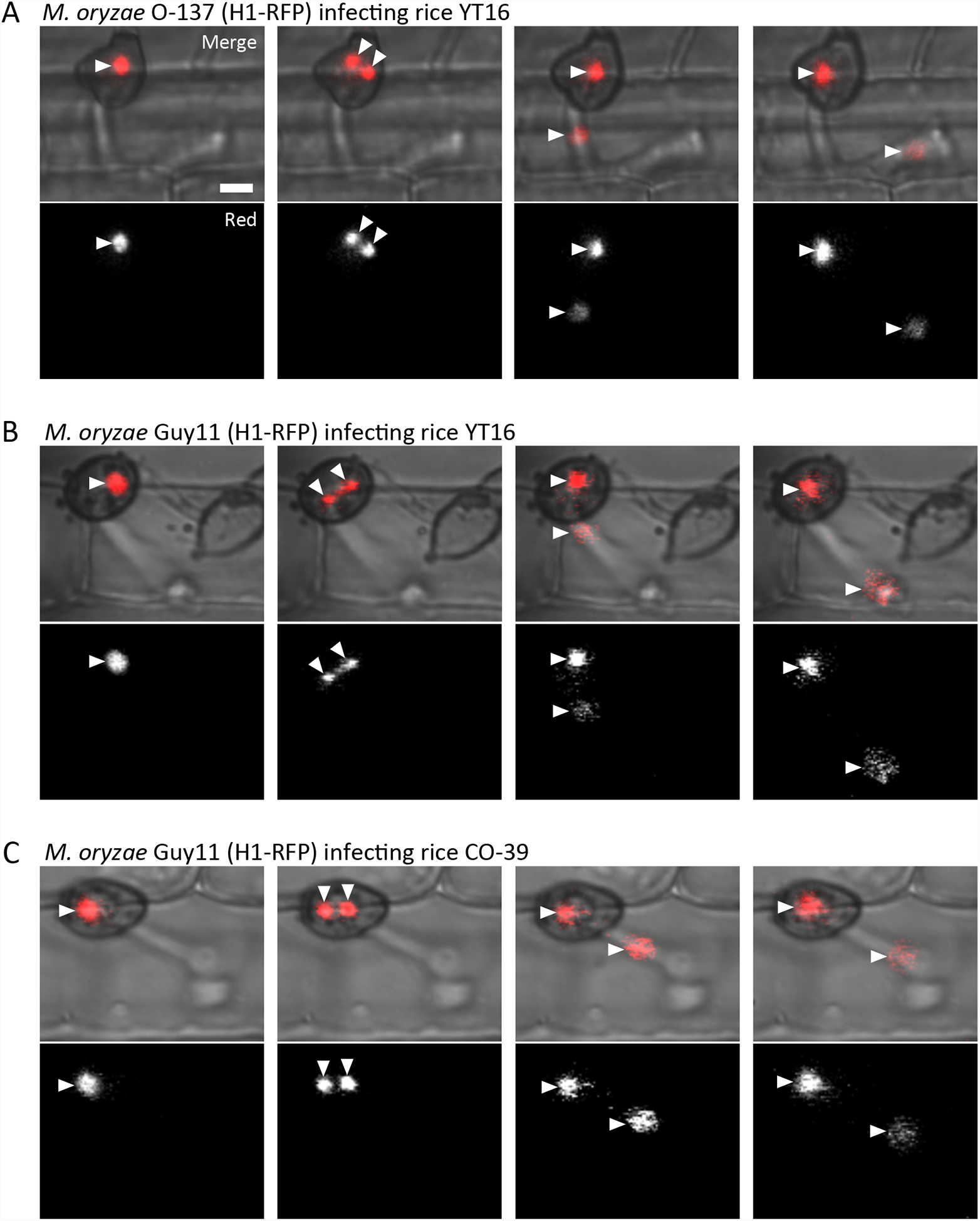
Nuclear division occurs within the appressorium. (A) *M. oryzae* CKF1962 (derived from O-137) infecting rice cultivar YT16. Shown are single plane confocal images of both red fluorescence and bright-field (top), and red fluorescence alone (shown in white; bottom). Nuclei are denoted by arrowheads. A single nucleus tagged with tdTomato (red) separated into two mitotic nuclei within the appressorium at 26 hpi. One nucleus then migrated out of focus into the primary hypha, while the other remained in the appressorium (B) *M. oryzae* Guy11 infecting YT16 at the same growth stage as (A). Nuclear dynamics were consistent with (A), with nuclear division occurring within the appressorium (double arrowheads) and migration of a single nucleus out of focus into the primary hypha.(C) *M. oryzae* Guy11 infecting rice cultivar CO-39 at the same growth stage as both (A) and (B). Nuclear dynamics were again consistent with nuclear division occurring in the appressorium (double arrowheads) and migration of one nucleus into the primary hypha. Bar = 5 µm.

### Formation of the first septum after nuclear division and migration

To define the location and timing of the septation during development of the first IH cell, we generated *M. oryzae* transformants expressing fluorescently labeled septa together with H1-tdTomato. The septa were visualized by constitutively expressing the plasma membrane targeted PLC delta:PH as a translational fusion to GFP (GFP-PLC delta:PH) in *M. oryzae.* PLC delta:PH binds to PI(4,5)P2 lipids [19] and has been shown to localize to the plasma membrane and septa in *N. crassa* [20]. *M. oryzae* transformants expressing GFP-PLC delta:PH showed green fluorescence localized at the plasma membrane of the appressorium and primary hypha (Fig. 3 top). After nuclear division, one nucleus moved to the bulbous tip of the primary hypha, and the other nucleus remained in the appressorium (Fig. 3 middle). Subsequently, the green fluorescence localized near the midpoint of the filamentous portion of the primary hypha, indicating the site of the first septum (Fig. 3 bottom). The primary hypha was previously characterized by Heath et al (1990) and Kankanala et al (2007): it is a filamentous structure that differentiates from the penetration peg, an extremely narrow structure that originates at the appressorium and forces through the plant cell wall, and that subsequently differentiates into bulbous IH. Our study further shows that the primary hypha is a filamentous part of two distinct cell types, constituting the invasive extension of the single-celled appressorium and the first IH cell.

**Figure 3.**
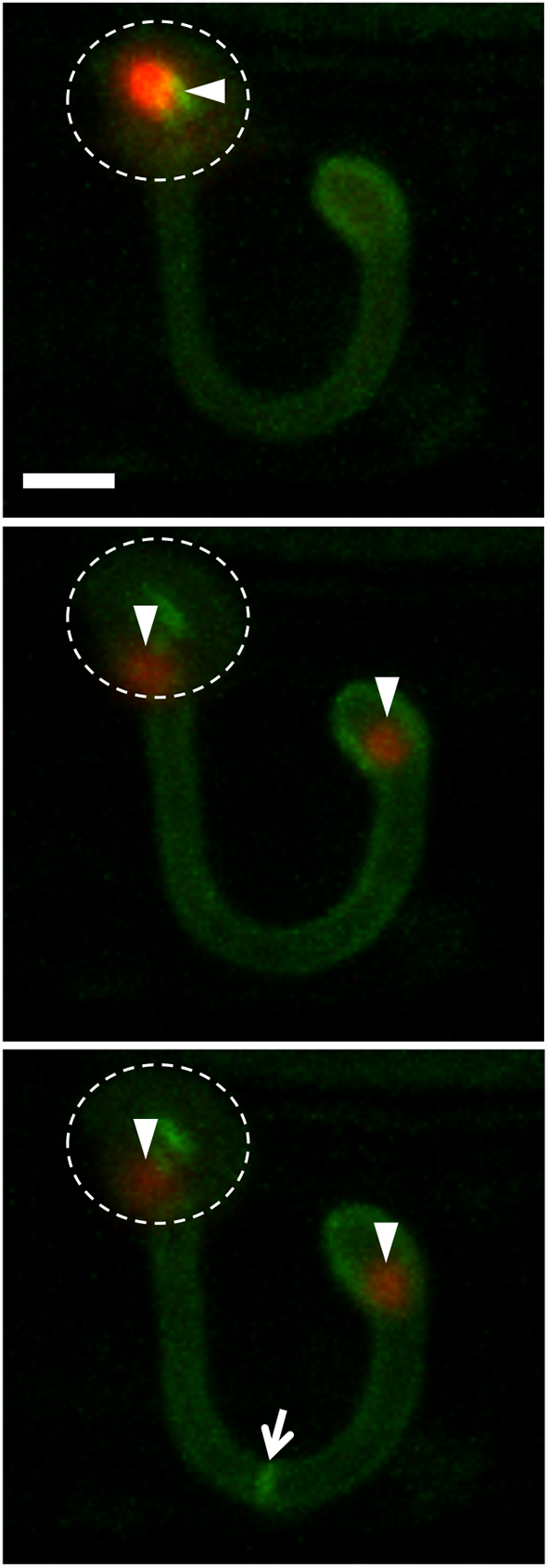
The first post-penetration septum forms in the primary hypha. Shown are maximum intensity projections of confocal optical sections of a rice cell invaded by *M. oryzae* strain CKF2686 at 27 hpi. Merged nuclear tdTomato (red) and GFP fused to PLCdelta:PH (green) are shown. Top: One nucleus (arrowhead) was located in the appressorium (outlined by dotted line) and GFP-PLCdelta:PH was localized to the plasma membrane. Middle: The same infection site imaged 50 minutes later showing one appressorial nucleus and one nucleus in the bulbous hypha. Bottom: The same infection site after an additional 24 minutes showing GFP fluorescence concentrating at the location of the first septum (arrow). Bar = 5 µm.

### Development of BICs during hyphal differentiation

Two-stage development of BICs has been previously documented [11] (Fig. 1A): the tip BIC is associated with the apex of the primary hypha; as the primary hypha switches to bulbous IH growth, the BIC maintains its position on what is now the side of the bulbous IH. To determine how the tip BIC becomes the side BIC, we used our earlier observations about the time of nuclear division relative to hyphal swelling (Fig. 1C and Fig. 2) to design an experiment. We imaged *M. oryzae* transformant CKF1651 expressing cytoplasmic EYFP and nuclear mRFP as well as mRFP fused to the BIC-localized cytoplasmic effector Pwl2 [11] at different growth stages analogous to the three panels in Fig 1B. In cases of no bulbous differentiation, the BIC was at the tip of the primary hypha (Fig. 4A left). When bulbous differentiation had occurred, but nuclear division had not, the BIC was already at the side, indicating that the BIC is displaced from its tip position as the primary hypha swells (Fig. 4A middle). In infections where nuclear division and migration had taken place, we observed the BIC still at the side, where it remained as a new hypha began to grow at the tip (Fig. 4A right). These results indicate that the bulbous swelling of the primary hypha occurs prior to nuclear division and is responsible for the BIC being “left” on the side of the cell. This displacement of the BIC indicates that there is depolarization at the apex of the primary hypha during bulbous differentiation. The position of the BIC on one side of the BIC-associated cell suggests that the opposite side of the primary hypha grows more than the BIC side. This asymmetric swelling produces an oval cell offset from the center of the hypha (Fig. 1 and Fig. 4) rather than a round cell, which would result from equal swelling in all directions. This growth is clearly distinct from the polarized growth observed in filamentous primary hypha development, of which the tip BIC is characteristic. Polarity in filamentous fungi is established and maintained by multiple proteins, especially those associated with control of cytoskeletal positioning [21]. The Spitzenkörper (Spk) and polarisome have been established as instrumental in determining the direction of hyphal tip growth in filamentous fungi [22, 23, 24]. In *M. oryzae*, Spk-associated motor protein Mlc1 is localized to the growing tip of the primary hypha before differentiation of the BIC-associated cell. After differentiation, Mlc1 is not present at the growing IH tip, but can be visualized near septa and in the subapical BIC-associated cell [17]. In contrast, the polarisome component Spa2 is localized to the growing tip of the primary hypha and IH [17]. Because an IH cell develops from the growing end of the BIC-associated cell, there must be a repolarization event after asymmetric swelling occurs. The Spa2 and other polarisome proteins are likely involved in repolarization of the IH growing tip.

**Figure 4.**
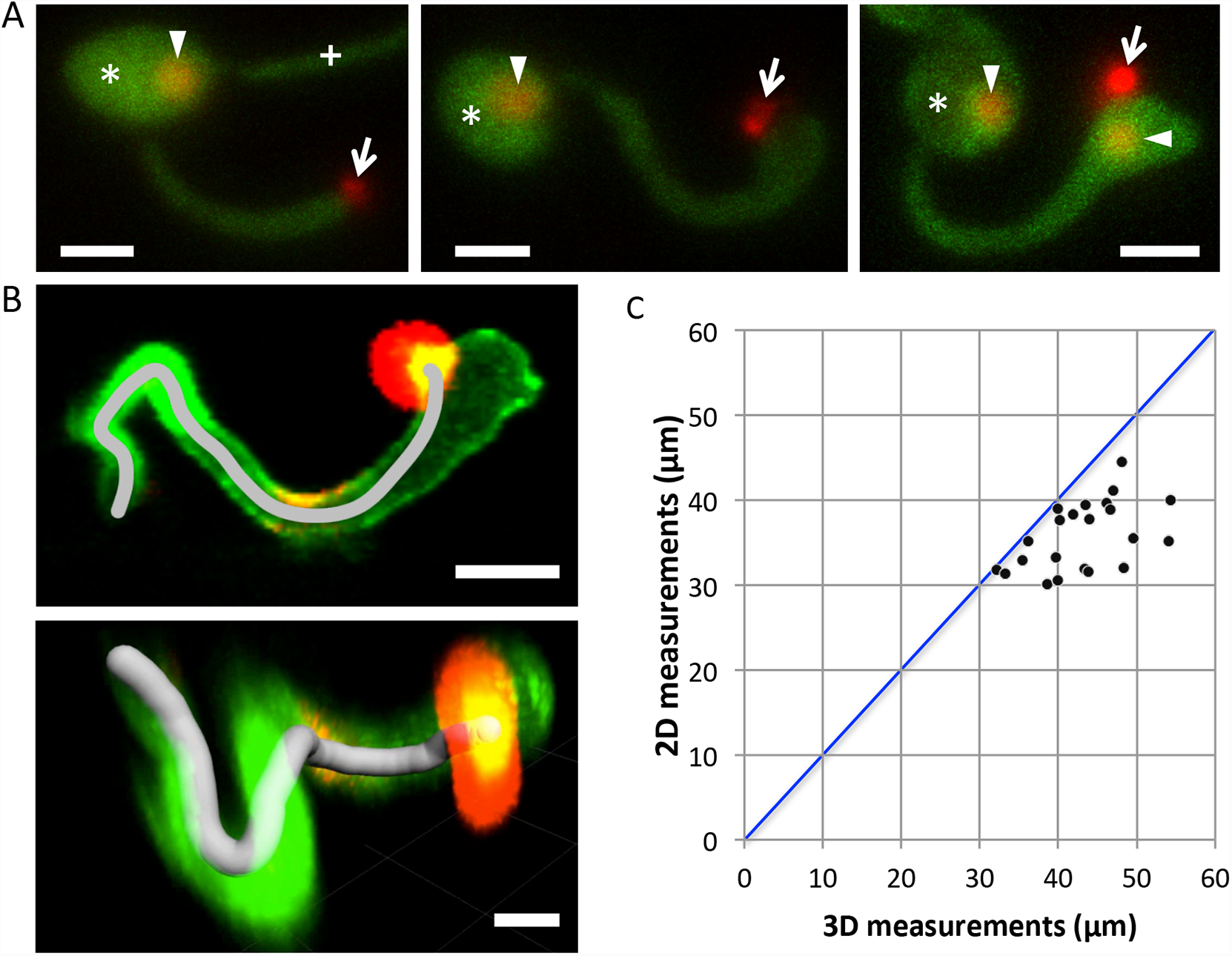
BIC development. (A) Tip to side BIC differentiation and first nuclear division event at 26 hpi. Shown are maximum intensity projections of confocal optical sections (18.9 - 28.5 µm in total depth). Images are of *M. oryzae* CKF1651, expressing cytoplasmic EYFP (green), nuclear mRFP (red), and BIC-localized mRFP (red; cytoplasmic effector Pwl2 fused to mRFP). Arrows = BICs. Arrowheads = nuclei. Asterisk = appressorium. Left: Tip BIC stage. Middle: Side BIC stage. Right: After nuclear division, the side BIC remained at the same position. Bars = 5 µm (B) Representative measurement from the penetration point to the side BIC in *M. oryzae* CKF1616 at 29 hpi. Images show BIC-localized mCherry (red; cytoplasmic effector Pwl2 fused to mCherry:NLS), fungal hypha outlined by EGFP (green; apoplastic effector Bas4 fused to EGFP), and mCherry and EGFP overlapping in the side BIC (yellow). Top: A maximum intensity projection of confocal optical sections (23 µm in total depth) and 2-dimensional length (gray line; length = 31.4 µm) measured with the open Bezier tool in the Zen Black software. Bottom: A 3-dimensional reconstruction of the same infection site shows 3D length (gray line; distance = 33.2 µm) measured with the Imaris dendrite creation algorithm (still image of the last frame of Supplemental Movie). Bars = 5 µm. (C) Comparative graphical summary of measurements taken with 3D Imaris and with 2D Zen Black software of the distance between the penetration point to the side BIC. Individual infection sites (black dots) have both a 3D measurement (x-axis) and a corresponding 2D measurement (y-axis). The vertical distance between a data point and the 1-to-1 line (blue) on the graph corresponds to the difference in length between the 2D and 3D measurements associated with that data point.

### Regular growth from the appressorium penetration to the side BIC

The filamentous primary hypha appeared to extend from the appressorium to a consistent distance before widening at its apical region. We reasoned that the side BIC would be positioned at a defined distance from the appressorium. To test this, we measured the distance from the appressorium penetration point to the side BIC using *M. oryzae* transformant CKF1616,expressing the BIC-localized cytoplasmic effector Pwl2 fused to mCherry:NLS and also IH-outlining apoplastic effector Bas4 fused to EGFP 11. The penetration point was identified by examining the EGFP fluorescence outlining the primary hypha in conjunction with the bright-field, and the BIC was identified based on the strongly localized mCherry fluorescence. We conducted two-dimensional measurements of these images using the Zen Black open Bezier tool (Version 8.1, Zeiss). The distance along the primary hypha from the penetration point to the side BIC has an average of 36 µm, ranging from 30 µm to 45 µm (n=22) (Fig. 4B top; Fig. 4C). To obtain more accurate measurements of the same infection sites, we also used the three-dimensional dendrite measurement algorithm in Imaris (Version 7.6, Bitplane). An additional movie file shows the three-dimensional measurement in more detail (see Additional file 1). The average distance was 43 µm, ranging from 32 µm to 54 µm (n=22) (Fig. 4B bottom; Fig. 4C; Additional file 1). Our finding of this considerable distance raises an intriguing question concerning effector secretion. Cytoplasmic effectors such as Pwl2 are secreted into the tip BIC and the side BIC, which are located more than 32 µm from the appressorial nucleus (Fig. 4A left and middle; Fig. 4B-C) in which the *PWL2* gene is expressed (Zhu and Khang, unpublished). There must be a mechanism to transport effectors from the appressorium through the primary hypha for their secretion to the distantly located BIC, but the mechanism remains to be determined.

### Hyphal growth inside heat-killed rice cells

We hypothesized that the development we observed during the early biotrophic invasion is specific to live host cells. To test this, we used a heat-killed cell assay, inoculating nonviable rice tissue with conidia. When inoculated on this tissue, *M. oryzae* produced the normal appressorium (Fig. 5), indicating that the surface cues were still sufficient for appressorium differentiation. After penetration into heat-killed host cells, the fungus did not show distinct differentiation from filamentous to bulbous growth as observed during invasion of live host cells, but elongated, filamentous-like growth throughout the rice cell and its neighbors (Fig. 5). Hyphae growing in heat-killed cells spread rapidly into adjacent cells rather than completely filling the first cell, which the IH in living cells do over the course of 12 hours [4]. In living host cells, BICs are easily recognized in bright-field images due to their predictable locations on the tip of the primary hypha, the side of the first bulbous IH cell, and the tips and sides of hyphae moving into adjacent host cells [11] (Fig. 1A). We did not observe any indications of BICs in the bright-field images of infections in heat-killed cells. The absence of BICs in dead host cells is consistent with Giraldo et al (2013) showing that BICs are derived from host membrane components such as the plasma membrane and endoplasmic reticulum; therefore, BICs are unlikely to form in host cells, which lack intact membranes.

**Figure 5.**
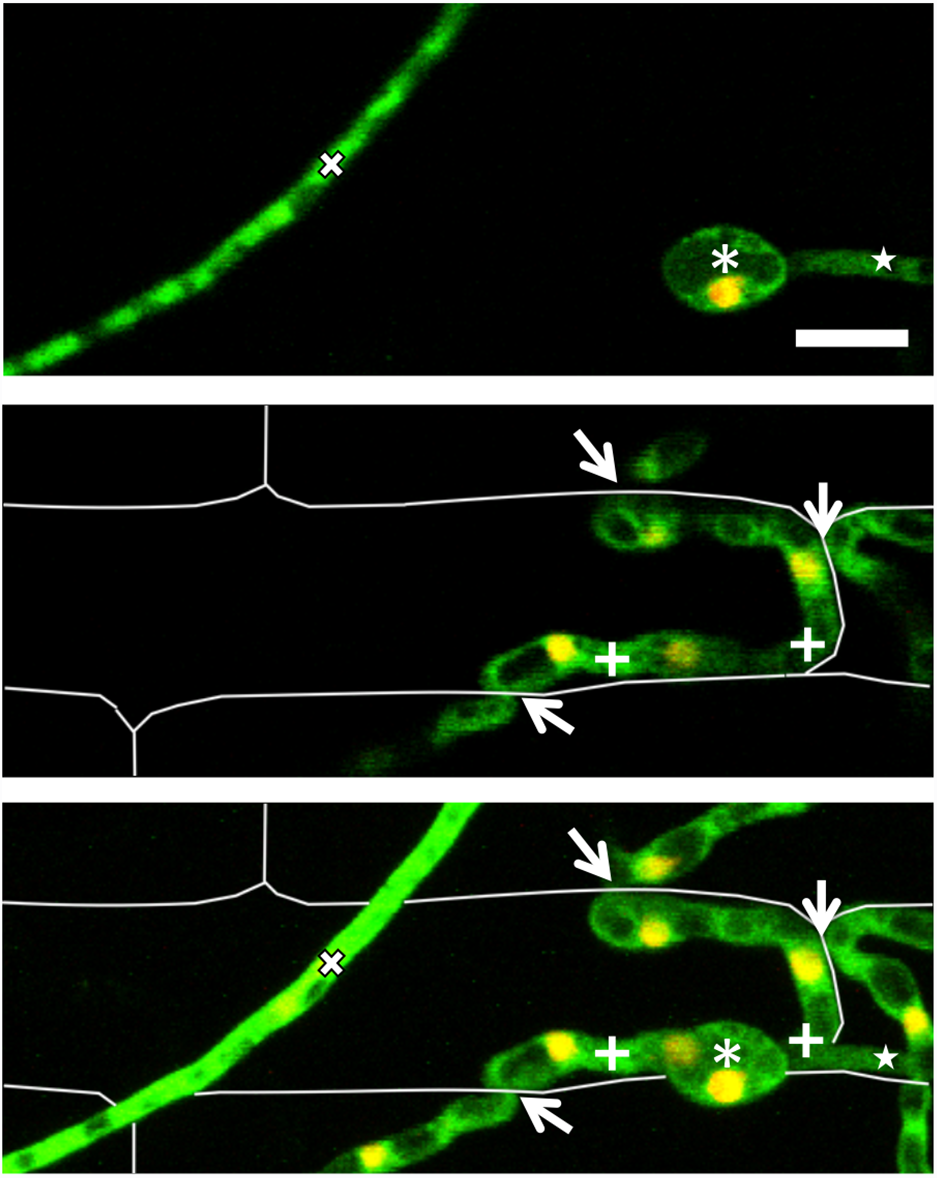
Hyphal morphology in heat-killed rice cells is distinct from growth in live cells. *M. oryzae* CKF2138, expressing cytoplasmic EYFP (green) and nuclear tdTomato (red, but shown in yellow due to overlap of green and red), growth in heat-killed rice cells at 25 hpi. Shown are merged fluorescence confocal images of a single focal plane (top and middle), or a projection (29 µm in total depth) (bottom) of the same infection site. Rice cell walls are shown as white lines. Top: A single focal plane shows the appressorium (asterisk) with a nucleus, germ tube (star), and hypha that grew on the rice surface (x). Middle: A focal plane 12 µm beneath the appressorium shows 2 filamentous-like IH branches (plus signs) and locations where IH had crossed the rice cell wall into neighboring cells (arrows). Bottom: A maximum intensity projection shows all features of both focal planes (top and middle). Bar = 10 µm.

## Conclusions

We provide cellular details of early post-penetration development of *M. oryzae* during invasion of susceptible rice cells. This study defines six developmental stages, in which the single-celled appressorium gives rise to the first IH cell, and the tip BIC becomes the side BIC when the tip BIC-associated hyphal apex undergoes asymmetric swelling prior to mitosis in the appressorium (Fig. 6). Development of BICs and BIC-associated cells is induced during invasion of live rice cells. Our studies provide a descriptive framework for early biotrophic invasion and lead to new mechanistic questions concerning hyphal differentiation in response to host environmental cues and effector trafficking from the appressorium to BICs.

**Figure 6.**
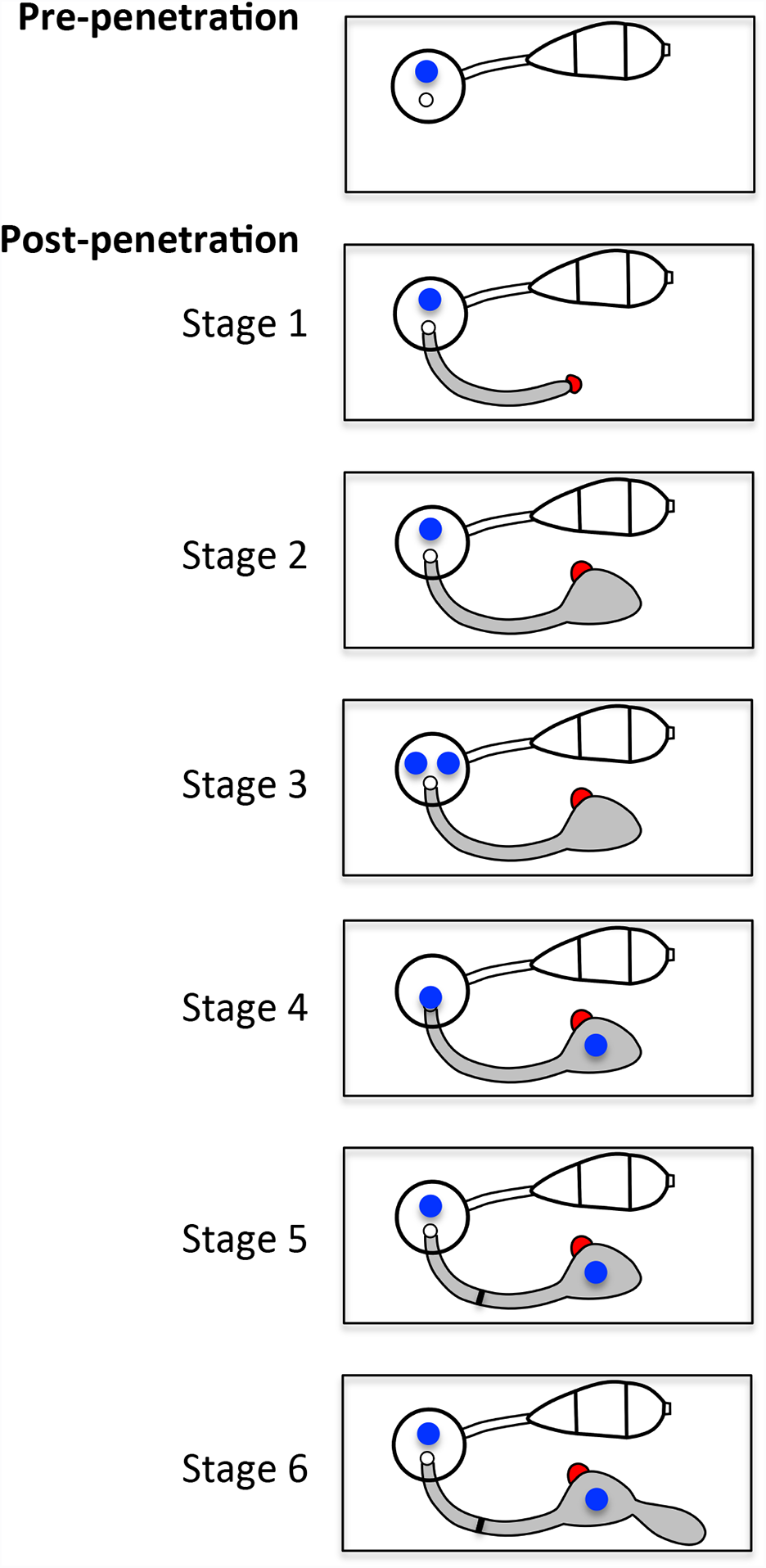
Schematic diagram of early post-penetration growth in susceptible rice cells. Pre-penetration: *M. oryzae* conidium (white; three-celled) germinates and forms an appressorium (circle) containing one nucleus (blue circle). Post-penetration: (Stage 1) The appressorium penetrates the host cell wall, forming an anucleate filamentous primary hypha (gray) with the tip BIC (red). (Stage 2) The tip of the filamentous primary hypha swells and the tip BIC becomes the side BIC (red). The primary hypha remains anucleate. (Stage 3) Nuclear division occurs in the appressorium. (Stage 4) One mitotic nucleus migrates into the bulbous portion of the primary hyphal tip, while the other nucleus remains in the appressorium. (Stage 5) The first septum forms in the filamentous portion of the primary hypha (blue line), finalizing the establishment of the effector-secreting BIC-associated cell. (Stage 6) Polarized growth resumes from the BIC-associated cell.

## Methods

### Plasmids, strains, and fungal transformation

For nuclear-localized tandem-dimer Tomato (tdTomato), the binary plasmid pCK1287 was constructed by cloning the histone H1 gene from *Neurospora crassa* (H1) at the 5’ end of the tdTomato gene under control of the *M. oryzae* ribosomal protein 27 (P27) promoter as follows: the 2.2 kb *Eco*RI-*Bam*HI fragment (P27:H1) isolated from pBV229 and the 1.7 kb *Bam*HI-*Hind*III fragment (tdTomato:Nos terminator) isolated from pBV359 were ligated between *Eco*RI and *Hind*III sites of the binary vector pBV1 (pBHt2) [25]. For the membrane-localized GFP, the binary plasmid pCK1417 was constructed by cloning the following three fragments between *Eco*RI-*Hind*III sites of pBV141 (pBGt) [26]: (a) the 0.5 kb *Eco*RI-*Nhe*I fragment (P27) isolated from pCK1374, (b) the 1.4 kb *Nhe*I-*Hpa*I fragment (GFP-PLCdelta) isolated from pGFP-C1-PLCdelta:PH [19] (Addgene plasmid # 21179), and (c) the 0.5 kb *Pvu*II-*Hind*III fragment of pBV210 (*N. crassa* β-tubulin terminator). To construct pCK1374, the 0.5 kb PCR product containing the P27 promoter was amplified from pBV167 using the primers CKP23 (5’-GAATTCGAATTGGGTACTCAAATTGG-3’) and CKP355 (5’-GCTAGCTTTGAAGATTGGGTTCCTAC-3’) and subsequently cloned in pJET1.2 (Thermo Scientific). The underlined sequences in CKP23 and CKP355 correspond to *Eco*RI and *Nhe*I sites, respectively. pBV plasmids (pBV1, pBV141, pBV167, pBV210, pBV229, and pBV359 as well as pBV377 described below) were obtained from Dr. Barbara Valent (Kansas State University).

Plasmids were transformed into *M. oryzae* O-137 or O-137-derived transformants using *Agrobacterium tumefaciens*-mediated transformation [27]. O-137 is a highly aggressive isolate collected from rice (*Oryza sativa*) in Hangzhou, Zhejiang, China [28]. CKF2138 (used in Fig. 1B, C, and Fig. 5) was made by transforming KV1 with pCK1287 (P27:H1-tdTomato). KV1 is an O-137-derived strain that constitutively expresses cytoplasmic EYFP (P27:EYFP) [4]. CKF1962 (used in Fig. 2A) was made by sequentially transforming O-137 with pCK1288 (P27:3xGFP:nuclear localization signal or NLS) and pCK1287 [8]. *M. oryzae* Guy11 (H1-RFP; used in Fig. 2B and 2C) has been described in Fernandez et al (2014). CKF2686 (used in Fig. 3) was made by sequentially transforming O-137 with pCK1287 and pCK1417 (P27:GFP-PLCdelta:PH). CKF1651 (used in Fig. 4A) was made by transforming the O-137-derived CKF110 (P27:EYFP and P27:H1-mRFP) with pBV377 (expressing cytoplasmic effector protein Pwl2 fused to mRFP with the native *PWL2* promoter) [11]. CKF1616 (alias KV121; used in Fig. 4B) is an O-137-derived strain that expresses cytoplasmic effector Pwl2 fused to mCherry:NLS with the native *PWL2* promoter and apoplastic effector Bas4 fused to EGFP with the native *BAS4* promoter [11]. Fungal transformants were purified by the isolation of single germinating spores. Wild-type strain O-137 and transformants were stored dehydrated and frozen at −20°C to maintain full pathogenicity and cultured on oatmeal agar plates at 24°C under continuous light [29].

### Infection assays

Rice sheath inoculations were performed as previously described [10, 30]. Briefly, excised leaf sheaths (5-9 cm long) from 17- to 21-day old plants were inoculated with a spore suspension (5 x 10^4^ spores/mL in sterile water). The inoculated sheaths were hand-trimmed at 22-28 hpi and immediately used for confocal microscopy. For heat-killed sheath inoculation assays, we first optimized heat-treatment conditions as follows: Trimmed sheaths were incubated in 70°C water for 10 or 25 min, immersed in 0.75 M sucrose, and then visually examined under a 20x objective lens for indications of plasmolysis. Sheath cells treated for 10 min underwent some plasmolysis, but those treated for 25 min did not, indicating the 25 min incubation was sufficient to kill cells. In subsequent heat-killed inoculations, pre-trimmed sheaths were incubated in 70°C water for 25 min, allowed to cool to room temperature, and then inoculated with a spore suspension.

### Microscopy and image analysis

Confocal microscopy was performed on a Zeiss Axio Imager M1 microscope equipped with a Zeiss LSM 510 META system using Plan-Apochromat 20X/0.8 NA and Plan-Neofluor 40x/1.3 NA (oil) objectives. Excitation/emission wavelengths were 488 nm/505 to 530 nm (EGFP and YFP), and 543 nm/560 to 615 nm (mRFP, mCherry, and tdTomato). Images were acquired and processed using LSM 510 software (Version 3.2). Confocal images of CKF1616 used for measuring the distance from the appressorial penetration point to the side BIC were obtained from Dr. Barbara Valent. For 2-dimensional measurements, confocal z-stack images were processed using the Zen Black software (Version 8.1, Zeiss). Each image was converted to a maximum intensity projection. The appressorial penetration point was identified based on brightfield visualization of the appressorium and on where the green fluorescence (Bas4-EGFP) outline of the primary hypha stopped. The BIC was identified from the overlap of red (Pwl2-mCherry:NLS) and green fluorescence. The distance between these two points was measured along the center of the primary hypha using the open Bezier tool. For 3-dimensional measurements, confocal z-stack images were imported into Imaris software (Version 7.6, Bitplane). The penetration point and BIC were identified as described for 2D measurements. The dendrite creation algorithm was used to produce a filament between the two points based on green fluorescence outline of the primary hypha for determining the path, and the length of the created dendrite was measured.

## Acknowledgements

We thank Drs. Seogchan Kang, Barbara Valent, and Richard Wilson for sharing materials. We thank current and former members of the Khang Lab (http://www.khanglab.org/) for support and discussions. We acknowledge the assistance of the Biomedical Microscopy Core at the University of Georgia with imaging using a Zeiss LSM 510 confocal microscope. This work was supported by the Agriculture and Food Research Initiative competitive grants program, Award number 2014-67013-21717 from the USDA National Institute of Food and Agriculture.

## Additional file 1

**Three-dimensional measurement from the appressorium penetration point to the BIC of.*M. oryzae* strain CKF1616 growing in a rice cell at 29 hpi.** Apoplastic effector Bas4 with a translational fusion to EGFP (green) outlined the invasive hypha. The BIC and the rice nucleus were visualized by BIC-accumulating cytoplasmic effector Pwl2 with a translational fusion to mCherry:NLS (red). Overlap of green and red signals is shown in yellow. Source image is a series of 1 µm confocal optical sections taken over a depth of 23 µm. Movie was created using the key frame animation feature in Imaris software (Version 7.6, Bitplane). At the start of the animation, the penetration point is on the left, and the BIC is on the right. As the animation rotates, a penetration point-to-BIC measurement (gray line) is overlaid on the image. The measurement was performed with the dendrite creation algorithm in the Imaris software (described in the Materials and Methods section) and has a length of 33.2 µm. A maximum intensity projection of the same infection site in this movie is shown in Figure 4B (top), and a mirrored still of the last frame in this movie is shown in Figure 4B (bottom). Bar = 5 µm

